# Filling gaps: fishing, genetics, and conservation of groupers, especially the comb grouper (badejo) (*Mycteroperca acutirostris*), in SE Brazil (2013-2020)

**DOI:** 10.1101/2021.04.12.439484

**Authors:** Alpina Begossi, Svetlana Salivonchyk, Branko Glamuzina, Alessandro Alves-Pereira, Carlos Eduardo De Araújo Batista, Regina H. G. Priolli

## Abstract

There are large gaps in our knowledge of the biology of important fish consumed by people in tropical countries, which makes conservation difficult. Small-scale fisheries are difficult to study and regulate, especially in countries with no systematic species monitoring. It is even more difficult to estimate the influence of these fisheries on vulnerable fish species and to diagnose possible damage to local fish populations. In this study, 490 individuals of **badejo**, or comb grouper (*Mycteropeca acutirostris)*, were observed at the Posto 6 fishery in Copacabana, Rio de Janeiro, for the periods of 2013-2014 and 2018-2020. A pattern of decreasing catches was observed for comb grouper. Therefore, provided that the fishing gear and the number of fish have remained the same, the apparent decrease in comb grouper needs to be further investigated. The results provide information regarding the reproduction of comb grouper, with major spawning season around spring (September-December) and additional spawning during April in SE Brazil. Samples from 96 groupers along the coast of Brazil were obtained, and genetic analyses were conducted. The genetic information obtained for grouper species enabled us to determine the relative genetic proximity of *M. acutirostris* and *Mycteroperca bonaci* and to obtain information that can be useful for aquaculture and conservation.

## Introduction

Small-scale fisheries (SSFs) are ubiquitous along maritime coasts. Despite a significant lack of systematic data, such fisheries are a valuable food source for local populations and are an important source of income [1,2]. Small-scale fisheries account for approximately 50% all fish captured for consumption worldwide; however, the lack of long-term monitoring in data-poor countries contributes to failures in fishery management [3].

Estimation of grouper populations and their distributions in coastal aquatic ecosystems is complex, as exemplified in a study on dusky grouper [4]. Of course, even small-scale fisheries can impact aquatic fauna when considering late-maturing species and the reality that environmental problems are often not considered to be of primary importance by local authorities, as is the case in Brazil [5]. On the other hand, it is known that SSFs can generally be sustainable even in developing countries (see examples of Caribbean and Latin American SSFs, such as those given by Salas et al. [6]). Therefore, the question is how are endangered, slowly growing species affected?

### The genus *Mycteroperca* and the comb grouper (*Mycteroperca acutirostris*)

In this group are the ‘badejo’, as they are called in Brazil. Their dorsal fins have 11 spines and 15-18 rays. They are coastal fish, have high commercial value, and are considered to be “noble fish”. Groupers are part of the Epinephelinae subfamily. Nevertheless, there is debate based on genetic data regarding whether the subfamily Epinephelinae should be treated as a family (Epinephelidae) and not a subfamily [7].

*Mycteroperca acutirostris* has a dark brown color and a head with long striations, and it is characterized by 11 dorsal spines and 15-17 rays and 50-56 rackers in the first branchial arch, along with a rounded caudal fin. It is associated with rocky bottoms (adults), and juveniles are found in shallow waters and mangrove areas; its major threat is fishing pressure [7,8]. Our studies have shown that this species spawns in the spring in Brazil [9,10].

Other species are addressed here, specifically in the genetic study, including *Mycteroperca bonaci*, called the black grouper, which has a dark brown color and regular, hexagonal spots; it has, among other traits, truncated caudal fins and 11-16 gill rackers in the first branchial arch. It is a reef species, but juveniles can be found in estuarine environments [7,8]. Another species included in the genetic study is *Mycteroperca insterstitialis*, called the yellowmouth grouper, which has small brown spots, 11 dorsal fins, 16-18 rays, a first branchial arch with 15-19 gill rackers, and an emarginate caudal fin. It is also a reef species. We considered other species for the genetic study, especially because they are found in the coast of Brazil: *M. acutirostris* is common in the SE whereas *M. bonaci* is common in NE Brazil. *M. interstitialis* is found in both regions, among others.

These species are monandric hermaphrodites, and *M. bonaci* forms spawning aggregations. Fishing pressure is one of the main threats to these species; adults occupy reef habitats, and juveniles are found in estuarine and mangrove environments. Adults feed on fish [7].

### The lack of data

When searching in FishBase [11] and the IUCN Red List (data from 2016 for these fish) for *Mycteroperca acutirostris* (comb grouper), we found that the knowledge about this species had several gaps. There are gaps for *M. acutirostris*, which include important information, such as diet and reproduction. The available information is general (and is based on few studies) such as the following: It has high vulnerability and high prices; there is no information regarding eggs, spawning processes, or periods; and food items are mentioned as “unidentified invertebrates” (Froese and Pauly [11]: February 17, 2021: 14:35). Nevertheless, it is considered to be a species of LC (least concern), and its population is estimated as stable (UICN [12]: last assessed November 20, 2016). For *Mycteroperca bonaci*, for example, we found slightly more biological information (Froese and Pauly[11]: February 18, 2021: 11:38): spawning in Brazil (one study including other fish species) in the period of June-December occurs for *M. bonaci* [13], and its diet was unidentified. *M. bonaci* is considered to be NT (near threatened, population decreasing) (IUCN [14]: February 18: 11:42). For *Mycteroperca interstitialis* (yellowmouth grouper), we found some information regarding reproduction (a study in the United States) showing that its reproduction occurs in all months of the year and its diet consists of “nekton” (a study in Puerto Rico) (Froese and Pauly [11]: February 18: 11:52). It is considered to be VU (vulnerable), and its population is decreasing (IUCN [14]: February 18: 11:55).

It is important to mention here that even though we included some genetic data for other species of *Mycteroperca* in this study, which allows us to make comparisons, our main focus will be *M. acutirostris* because we have been following its catches for two periods since 2013 in Copacabana, Rio de Janeiro (State of Rio de Janeiro), with some scattered observations for Bertioga and Santos (State of São Paulo).

The objective of this study was to obtain data about the fishing activity and biology of the comb grouper. Given the substantial gaps in knowledge about an important food species and for the market, we believe that original and previously unpublished data regarding *M. acutirostris* are important for conserving this species.

### Our earlier studies

This study follows a series of studies on small-scale fishing for groupers, especially *Epinephelus marginatus* (dusky grouper) and *Mycteroperca acutirostris* (comb grouper), that began in 2006 for both the dusky grouper [15] and comb grouper [10] at the Colônia de Pescadores do Posto 6 in Copacabana, Rio de Janeiro, Brazil.

### Studies of other Epinephelinae: Epinephelus marginatus, dusky grouper

*Epinephelus marginatus* is found in the Atlantic and in the western Indian Ocean with a decreasing population trend and is considered by the IUCN to be “EN” (endangered); *Mycteroperca acutirostris* is found in the western Atlantic and is considered to be heavily fished, but it is viewed by the IUCN to be of “LC” (least concern) [7,11].

Here, we chronologically describe the research results for garoupa, dusky grouper. The most recent study [16] showed relative catches and price stability. The studies that were begun by Begossi and Silvano [15] included results about the local knowledge of fishers regarding diet and habitat, fishing locations, weight and length (TL), and stomach contents, among other features (sample of 40 individuals from 2006-2007). This study was followed by that of Begossi et al. [17], which included fishing locations (fishing spots), weight and length (TL), macroscopic analysis of gonads by trained fishers (800 individuals from 2013 to 2015); Begossi et al. [18]: locations, weight and length (TL), macroscopic analysis of gonads by trained fishers (sample of 222 individuals, from 2016-2017), and prices from October, 2016 to November, 2017; and work by Begossi and Salivonchyk [16]: 1,896 groupers from 2013-2018, with catches and prices being relatively stable in Copacabana (the https://www.biorxiv.org/content/10.1101/759357v1).

## Methods

### The fishery

The “Colônia de Pescadores do Posto 6” includes a small-scale fishing community on Copacabana Beach that was established in 1923. Fishing is conducted from small motorboats by using set gillnets, hooks and lines and by spearfishing [18,19]. Recently, spearfishing by diving has become important, especially among young fishers. Fishing effort has been similar over the last 11 years: there are approximately 20-25 active fishers, including approximately 10 divers who spearfish close to islands.

Fieldwork at Copacabana was undertaken from September 2013 to February 2020 at the landing point or fish market of Posto 6, Copacabana. One of the authors (AB) performed visits on approximately 5 days/month and compiled information on weights, prices, and locations, among other data. Two fishers were trained by following a protocol that is explained in detail in Begossi [20] and was used in Begossi et al. [17,18,21]. These methods included local knowledge, tracking fishery landings (systematic visits 3-5 days/month), including fishing locations, weight and length measurements (TL), and reproduction (gonad macroscopic analysis), which were the same as in the earlier studies [detailed in 18,20]. Garoupa, or dusky grouper (*E. marginatus*), was observed in 2013-2018, and comb grouper (*M. acutirostris*), was observed in two periods, namely, 2013-2014 and 2018-2020.

### Genetics of groupers

#### Sampling of fins for genetic analyses

A total of 96 samples were obtained along the coast of Brazil (see details in [18]) from specimens belonging to two genera (*Epinephelus* and *Mycteroperca*) and five grouper species: *E. marginatus* (N = 28), *E. morio* (N = 19), *M. acutirostris* (N = 16*), M. bonaci* (N = 27), and *M. interstitialis* (N = 6). All individuals were caught by commercial fishers and identified based on their morphological characteristics as described in Begossi et al. [18].

#### Molecular techniques

Total genomic DNA was extracted from approximately 20 mg of tissue using a DNeasy Blood and Tissue Kit (Qiagen, Hilden, GE). DNA concentrations were estimated using a Qubit v4.0 fluorometer (Thermo Fisher Scientific, Waltham, USA). Thermo Fisher Scientific) and were normalized to 20 ng/μl. Genomic libraries were constructed according to the Genotype-by-Sequencing double digestion protocol described by Poland et al. [22], using the restriction enzymes NsiI (NEB, Ipswich, USA) and MseI (NEB). The resulting libraries were pooled at 96-plex and sequenced on the Illumina NextSeq 500 sequencing platform (Illumina, Inc, USA) at the Hemocentro of Ribeirão Preto facilities (Brazil), in mid-output mode and set to produce 150 bp single-end reads. The quality of the obtained raw reads was assessed using FastQC software (http://www.bioinformatics.babraham.ac.uk/projects/fastqc/) at the Hemocentro of Ribeirão Preto facilities (Brazil).

For each genus, samples were demultiplexed, and the raw read sequences were filtered with the module “process_radtags” in the Stacks program (version 1.42) [23]. SNP calling retained only SNP per sequenced tag, with a minimum sequencing depth of 5X, frequency of the least common allele ≥ 0.05 and occurring in at least 90% of individuals within each species. The SNP identification was performed considering each genus separately, and also for the five species simultaneously. Population genomic analyses were performed by applying additional filtering parameters to obtain the maximum-quality SNPs: 1) individual samples with >55% missing data were excluded, and 2) SNPs with missing data in 25% of the samples or a minor allele frequency (MAF) <0.05 were removed.

#### Population genetic analyses

The filtered data were imported as a genind object into R and were analyzed mainly by using several packages for population genetics. An outlier locus approach was taken using the R packages PCAdapt version 3.0.4 [24] and fsthet [25] and the software package SelEstim [26], and only the data that were identified as outliers by at least two of the three methods were considered as candidates for selection. The number of groups considered in fsthet and SelEstim analyses were three for *Mycteroperca* two for *Epinephelus* (corresponding to the number of species of each collection sampled). The outputs from these analyses were used to create a neutral locus dataset and an adaptive locus dataset for further analysis. Neighbor-joining trees were generated for both the neutral and outlier datasets using Nei’s genetic distance method. Analyses of population structure were performed using discriminant analysis of principal components (DAPC) with the ‘Adegenet’ package [27].

## Results

### The fishery at Posto 6, Copacabana

The main fishing locations of the studied individuals of dusky grouper and comb grouper caught in 2013-2014 and 2018-2020 were the area in 2-7 km south from Copacabana beach, where a lot of archipelagos and islands are located: Cagarras Islands, Tijucas Islands, Redonda Islands, Rasa Island and others (Fig. 1).

**Fig 1.**
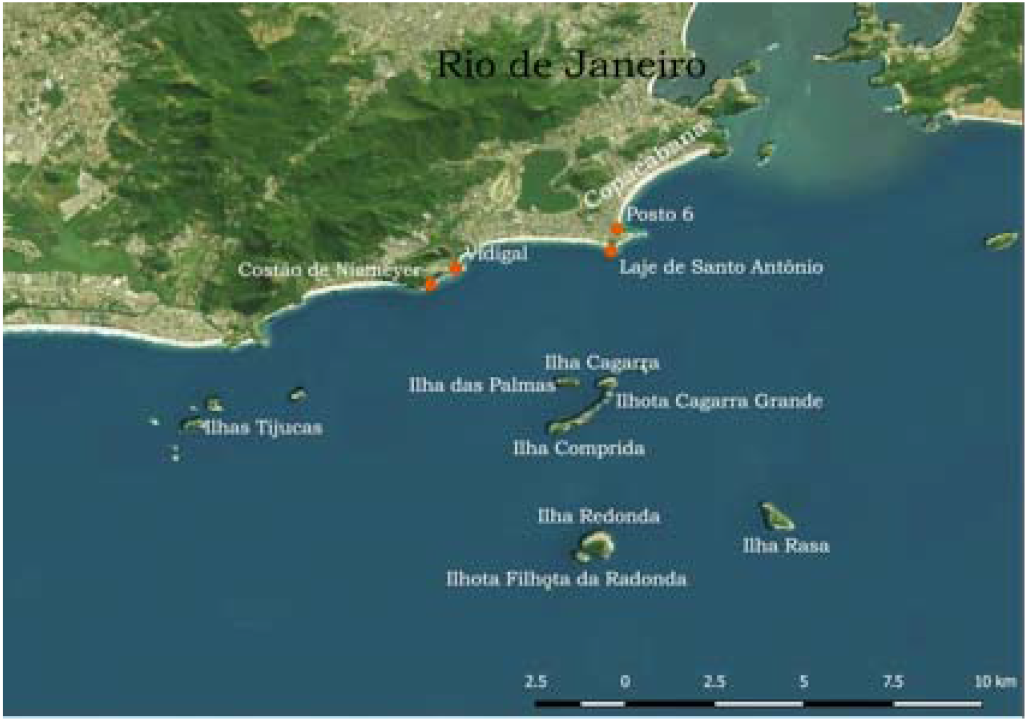
Fishing Locations Used for dusky grouper and comb grouper (2013-2018).

The number of individuals and the weights of comb grouper found and studied at the landing point and fish market in Posto 6, Copacabana significantly varied by year and month of the year (Tables 1 and 2).

**Table 1.** Comb groupers observed at Copacabana fishery from 2013 to 2021. The data in parentheses indicate the number of individuals for which the weight was determined, when it was not possible to weight the fish caught. Total number of individuals is 490 and weight obtained for 464 individuals (\****covid*** period). In the ***covid*** period of the study (May, 2020 to March 2021), occasional visits were made, and two trained fishers contributed through *whatsapp*, when possible. In this period of 27 days of observations (19 visits and 8 *whatsapp* messages), 19 *M. acutirostris* were observed.

| Month | 2013 | 2014 | 2018 | 2019 | 2020 |
| --- | --- | --- | --- | --- | --- |
| Jan |  | 14 |  |  | 23 (3) |
| Feb |  | 32 | 1 |  | 10 (5) |
| Mar |  | 12 |  |  | 2 |
| Apr |  | 26 | 3 (2) |  | * |
| May |  | 116 |  |  | * |
| Jun |  | 28 |  |  | * |
| Jul |  | 46 | 13 |  | * |
| Aug |  | 20 | 29 |  | * |
| Sep | 6 |  | 17 |  | * |
| Out | 34 |  | 21 |  | * |
| Nov | 18 |  |  | 2 | * |
| Dec | 3 | 1 |  | 17 | * |
| Total | 61 | 291 | 84 (83) | 19 | 35 (10) |

**Table 2.**
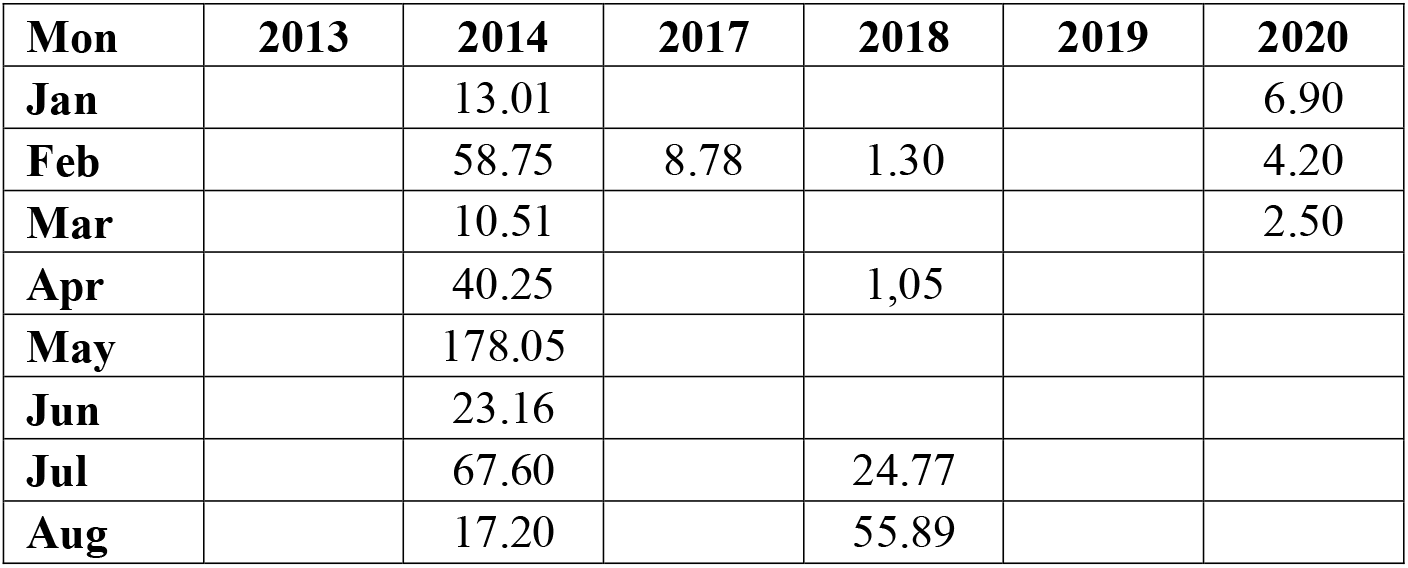
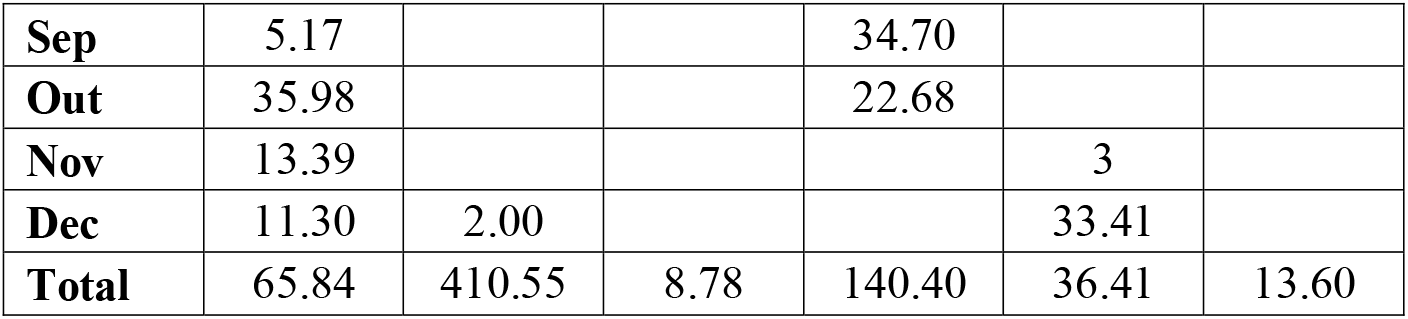
Weight (kg) of comb grouper caught at Copacabana fishery (2013-2020). Total weight is 630.39 (n=464 individuals) and average catch is 1.36kg. For the ***covid*** period (March 2020-March 2021), 27 days, the weight amounted 42,4 Kg (n=19), average 2,23Kg per individual.

| Mon | 2013 | 2014 | 2017 | 2018 | 2019 | 2020 |
| --- | --- | --- | --- | --- | --- | --- |
| Jan |  | 13.01 |  |  |  | 6.90 |
| Feb |  | 58.75 | 8.78 | 1.30 |  | 4.20 |
| Mar |  | 10.51 |  |  |  | 2.50 |
| Apr |  | 40.25 |  | 1,05 |  |  |
| May |  | 178.05 |  |  |  |  |
| Jun |  | 23.16 |  |  |  |  |
| Jul |  | 67.60 |  | 24.77 |  |  |
| Aug |  | 17.20 |  | 55.89 |  |  |
| **Sep** | 5.17 |  |  | 34.70 |  |  |
| **Out** | 35.98 |  |  | 22.68 |  |  |
| **Nov** | 13.39 |  |  |  | 3 |  |
| **Dec** | 11.30 | 2.00 |  |  | 33.41 |  |
| **Total** | 65.84 | 410.55 | 8.78 | 140.40 | 36.41 | 13.60 |

We observed a total of 490 individuals, among which weights were obtained for 466 individuals and showed a total catch of 630.39 kg. The average catch was 1.36 kg. While 410,55 kg was obtained in 2014, we observed a relative decrease in comb grouper catches, with 2019 being the year in which the species became rare in Copacabana fishing catches, and this trend continued in 2020.

Mature gonads for comb grouper, were observed in April, September and October: these spring months coincide with mature gonads being observed in dusky grouper [16,18,28] (Fig 2). The months mentioned by fishers from Copacabana (13 fishers) as the months in which comb grouper was “*ovado*” (with mature gonads) were September (5 fishers) and November-December (8 fishers).

**Fig 2.**
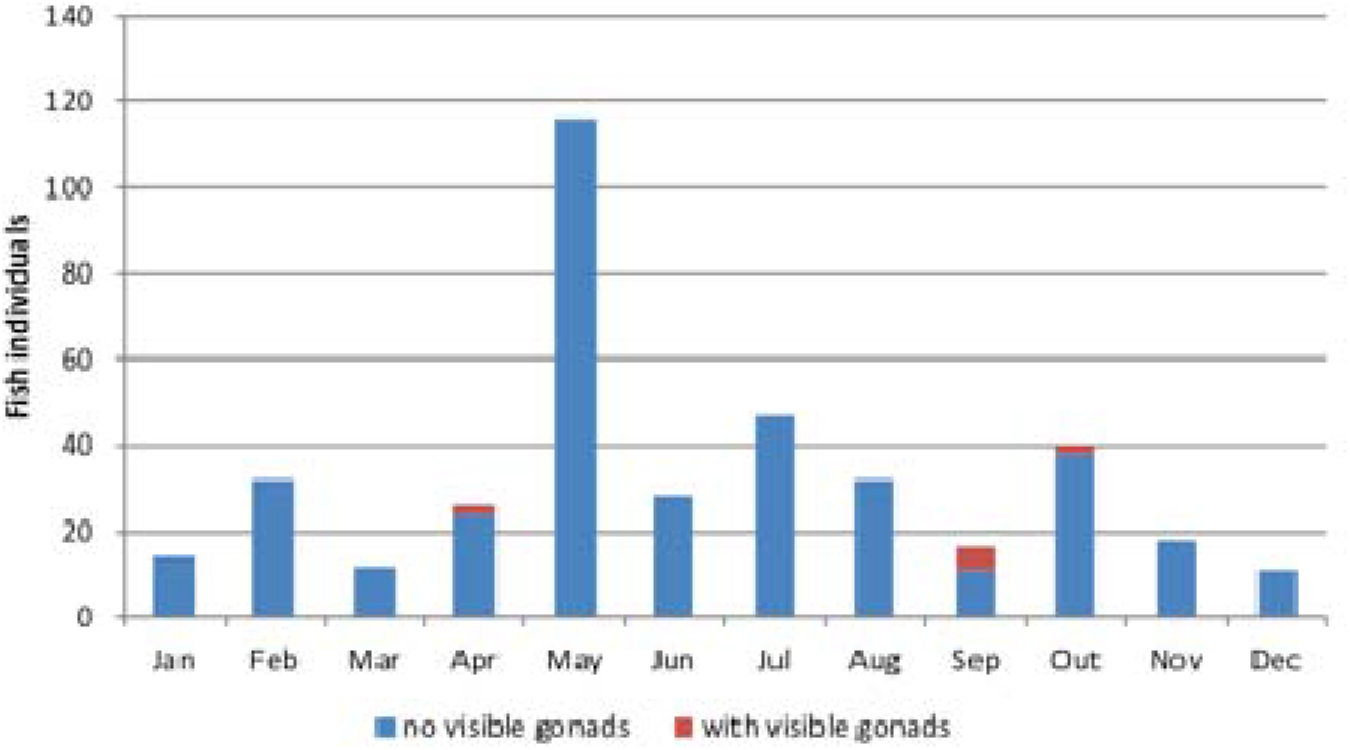
For the Period 2013-2020, 392 Individuals of Comb Grouper were Examined for Gonads. Visible Gonads were Observed in Only 9 Cases were: 5 in September and 2 in April and October.

Prices were examined for April and July October of 2018. The average price of comb grouper, was 40.8 Reais/kg. The prices ranged from 26.5 (26/07/2018) to 55.5 reais/kg (10/10/2018). The average daily prices varied from 36 (11/04/2018) to 45.4 reais/kg (10/10/2018) (the exchange rate on July 6, 2018, was 3.93 reais; Fig 2). The average monthly prices were slightly higher in July and August than in other months. They fluctuated most strongly in July and October (Figs 3a and b). The average exchange rate for 2018 was US$1.00=R$3.65) (https://www.exchangerates.org.uk/USD-BRL-spot-exchange-rates-history-2018.html).

**Fig 3a.**
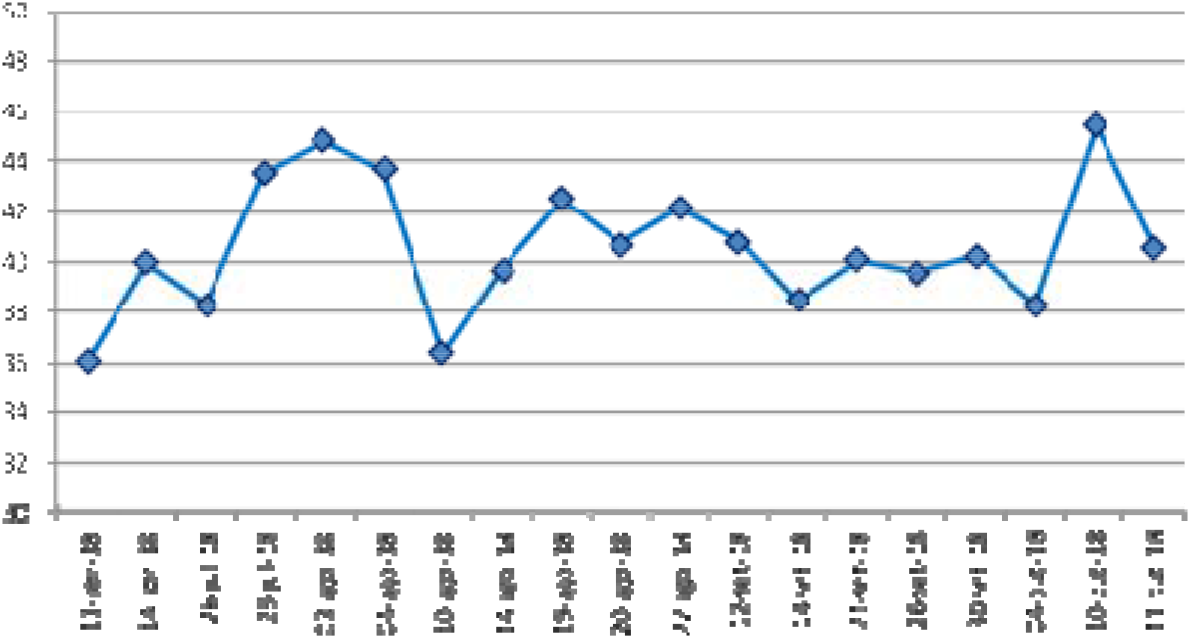
Daily Average Prices (in Brazilian Reais) per 1 kg of Comb Grouper, in Copacabana (2018, US$1.00=R$3.65).

**Fig 3b.**
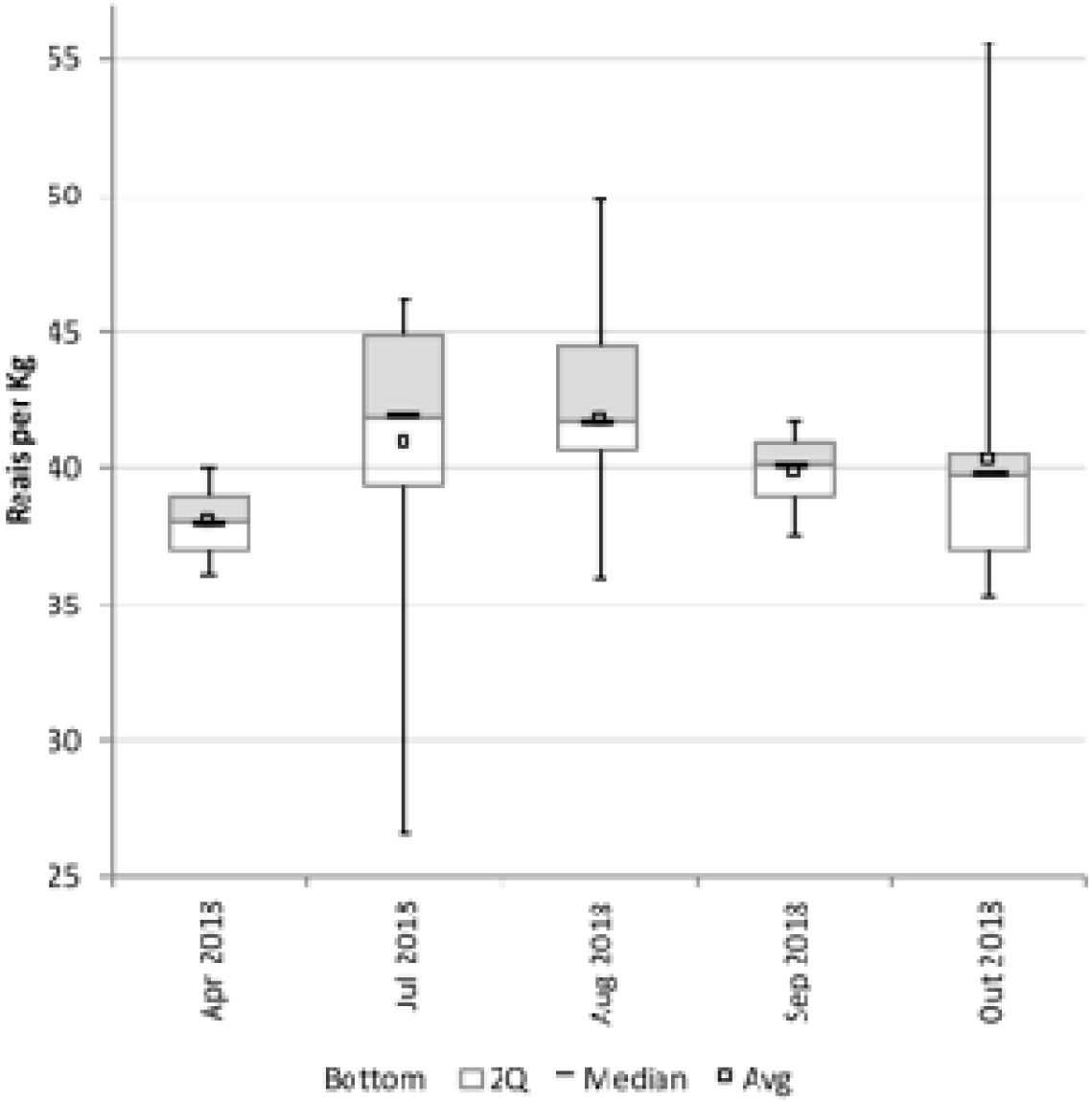
Monthly Average Prices (in Brazilian Reais) per 1 kg of Comb Grouper, in Copacabana (2018, US$1.00=R$3.65).

When comparing the yield (weight) of both species over the years, we observed higher catches for dusky grouper, compared to comb grouper, along with a decrease in comb groupers in 2018 and 2019 (Figure 4).

**Fig 4.**
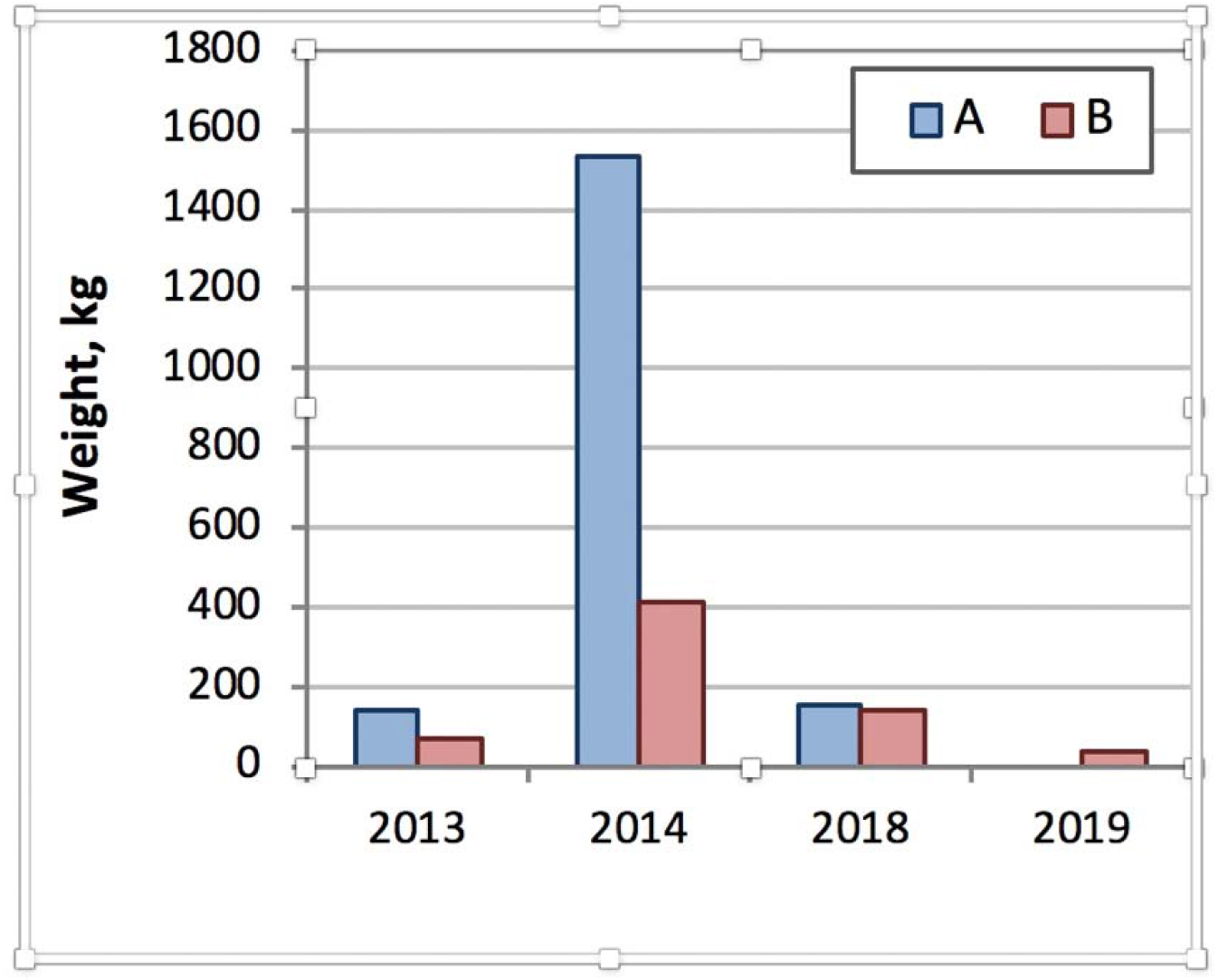
Annual Catches by Weight of Dusky Grouper (A) and Comb grouper (B). The results from samples taken from September 2013 to February 2018 for and from September 2013-December 2014 and January 2018-December 2019 for comb grouper at the Copacabana fishing post.

### Genetics of groupers in the coast of Brazil

GBS libraries produced 150,458,819 raw reads for the 96 groupers, and 139,429,799 reads were retained following “process_radtags” filtering. After applying the Stacks pipeline to each genus and additional filtering, 1313 SNP loci were identified in 43 individuals of *Mycteroperca* (*M. acutirostris* = 16, *M. bonaci* = 23 and *M. interstitialis* =4). For *Epinephelus*, 3528 SNP loci were identified in 45 individuals (*E. marginatus* = 27 and *E. morio* =18) (Figs A1 and A2, Appendix).

Outlier analyses identified 38 total consensus outlier loci in the three *Mycteroperca* species. Population structure analyses were conducted using the remaining 1,275 SNPs that were deemed to be neutrally evolving after outlier analyses. Overall, F_ST_ was 0.95 and highly significant (p < .001), while the species pairwise values varied from 0.935 to 0.967.

The high genetic divergence suggested by the pairwise F_ST_ estimates was also observed in the DAPCs. The analysis based on SNPs with neutral behavior explained almost 92% of the total variation in the first two components and clearly showed that samples of *M. acutirostris*, *M. bonaci* and *M. interstitialis* were highly distinct from each other (Fig 5). In Fig. 5, the DAPC scatter plot of *Mycteroperca acutirostris*, *M. bonaci*, and *M. interstitialis* that were collected in five Brazilian States (e.g., Rio de Janeiro, Bahia, Paraiba, Rio Grande do Norte), which were performed using 1,275 SNPs is shown.

**Fig 5.**
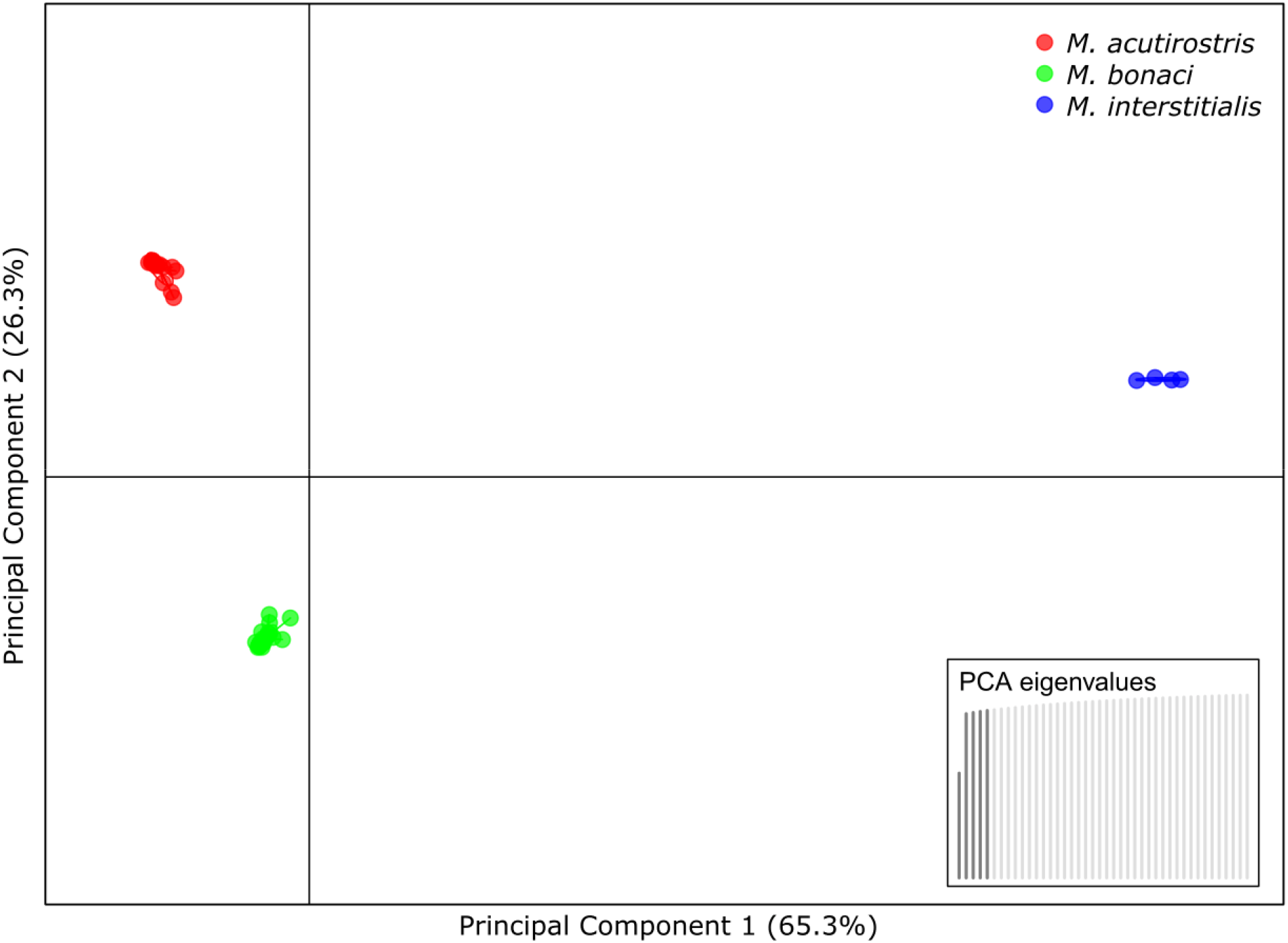
DAPC Plot of *Mycteroperca acutirostris*, *M. bonaci*, and *M. interstitialis*. Individual *Mycteroperca* species are represented by colored symbols, and the PCA eigenvalues show that the first two principal components explain more than 91% of the genetic variability.

The same clustering patterns were observed for the neighbor-joining trees using both the neutral and outlier datasets (Fig A1 Appendix), although individuals of *M. interstitialis* appeared to be less divergent to those of *M. acutirostris* than to those of *M. bonaci*.

The genetic distances between the five grouper species (Fig 6) were determined from a set of neutral SNP loci and revealed that *M. acutirostris* is more closely related to *M. interstitialis* (0.343) than to *M. bonaci* (0.48). *Epinephelus morio* was the species that was most distant from the others, except for another species from the same genus (*E. marginatus*) (Fig 6). In Fig. 6 neighbor-joining trees based on Nei’s genetic distances were produced using the set of 817 SNP loci identified with Stacks pipeline considering all the five Grouper species simultaneously.

**Fig 6.**
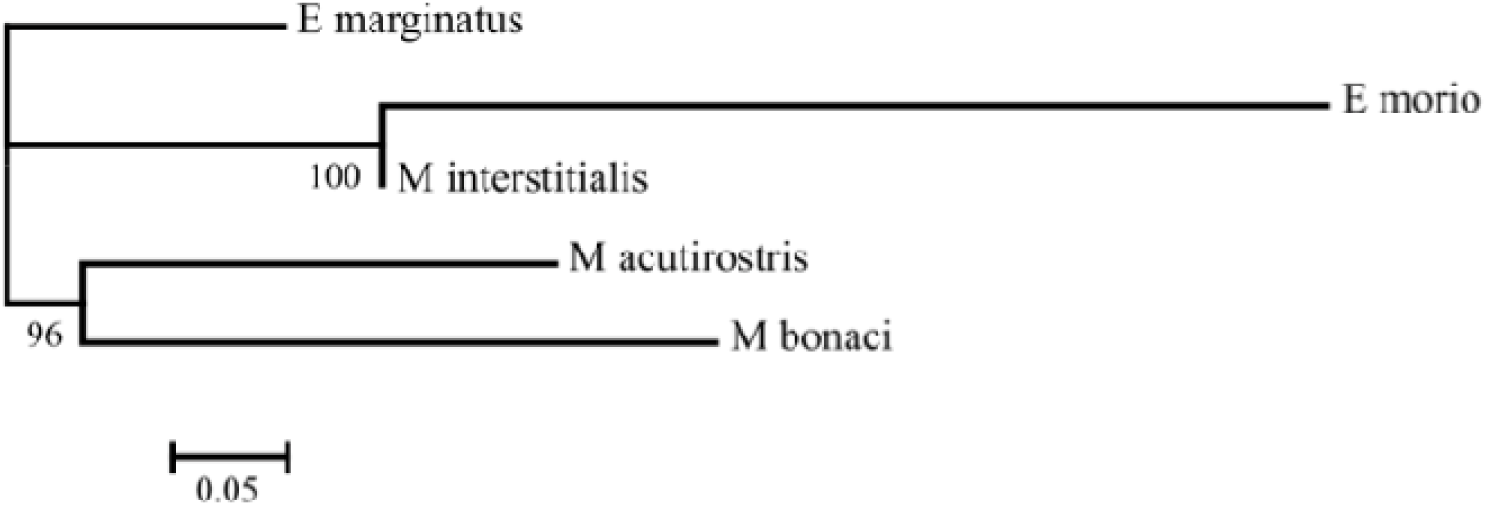
Neighbor-joining trees based on Nei’s genetic distances. Branch nodes are denoted as the percentage of bootstrap support that was generated with 1,000 replicates.

### Concluding remarks

Exploratory research is very important for species that require management, especially species that are important for human consumption and for the livelihoods of poor communities that depend on small-scale fisheries for consumption and cash. This study provides important information for the conservation of this species and provides information on the reproduction period and on genetics that may be useful for managing this species. In addition, it provides alerts for a species that is becoming scarce and for a species with poor data and that is still considered of LC (least concern).

In this study, we found different catch patterns for each of the grouper species, namely, stability of catches and prices for dusky grouper, and a decreasing pattern for comb grouper, for the Copacabana small-scale fishery. We added genetics to better understand the species and to gain insights into their conservation and biology. With this study, we hope to fill some gaps in Brazil and contribute to management efforts.

By comparing catches of dusky grouper and comb grouper, contrasting patterns are found in the catches of the studied groupers: while we found relative stability for the catches of dusky grouper, we found very irregular and decreasing catches in terms of number and weight for comb groupers. Even considering that comb grouper is of LC (least concern, IUCN Red List), there is an information gap concerning its diet and reproduction [11]. Thus, an optimistic prognosis is offered from the studies of dusky grouper and Copacabana [16], but there is a pessimistic prognosis for comb grouper due to its decreasing catches when the fishing effort is maintained at a constant level, as was observed for the Copacabana fishery.

The results for the genetics of groupers reveal that *E. morio* shows the greatest genetic distance from *Epinephelus*; however, the *Mycteroperca* subfamily has a relatively short genetic distance.

How could these data aid in the conservation of species? Such information can help directly and indirectly.

Directly, information regarding the reproduction period is helpful for managing fish resources, as fishing closures for comb grouper in the spring could help maintain stocks. Information on diet, in addition to its importance in maintaining available prey, could provide insights for possible aquaculture, which exists in SE Brazil for dusky grouper [29,30].

Indirectly, grouper genetics provides basic information for aquaculture; it engenders information that could aid in conservation by finding bottlenecks at some points, as was found for dusky grouper [31]. Here, genetic information could also be helpful for taxonomic analysis of the subfamily *Epinephelus*. Grouper systematics has been reevaluated, restricting the genus *Epinephelus* and expanding *Mycteroperca* when several species of Epinephelus were included in Mycteroperca, including *E. marginatus* [7,32].

## Acknowledgments

We are grateful to the fishers Antonio, Elenilson (“Jaguriçá”) and Raul; Ana helped with data from the Marimbás Club, Copacabana. AA-P thanks FAPESP (# 2018/00036-9) for a post-doctoral scholarship. AB is grateful to the productivity scholarship from CNPq (# 301592/2017-9), for the grant FAPESP # 14/16939-7, and for IDRC/UNICAMP grant (2019-2014). AA-P thanks FAPESP (# 2018/00036-9) for a post-doctoral scholarship. We are very grateful to Alline A. L. Tribst for the support in the laboratory, at Nepa/Unicamp, in the material extraction of the fish.

## Compliance with ethical standards

This research was approved and signed by B. R. Martins dos Santos, Comitê de Ética, Universidade Santa Cecília, number 1.747.889 on September 27, 2016 (Plataforma Brasil). It is approved under number 53824 at the SISBIO and is registered under number AB53669 at the SISGEN, MMA (Ministério do Meio Ambiente, Brasil).

## Appendix

**Fig A1.**
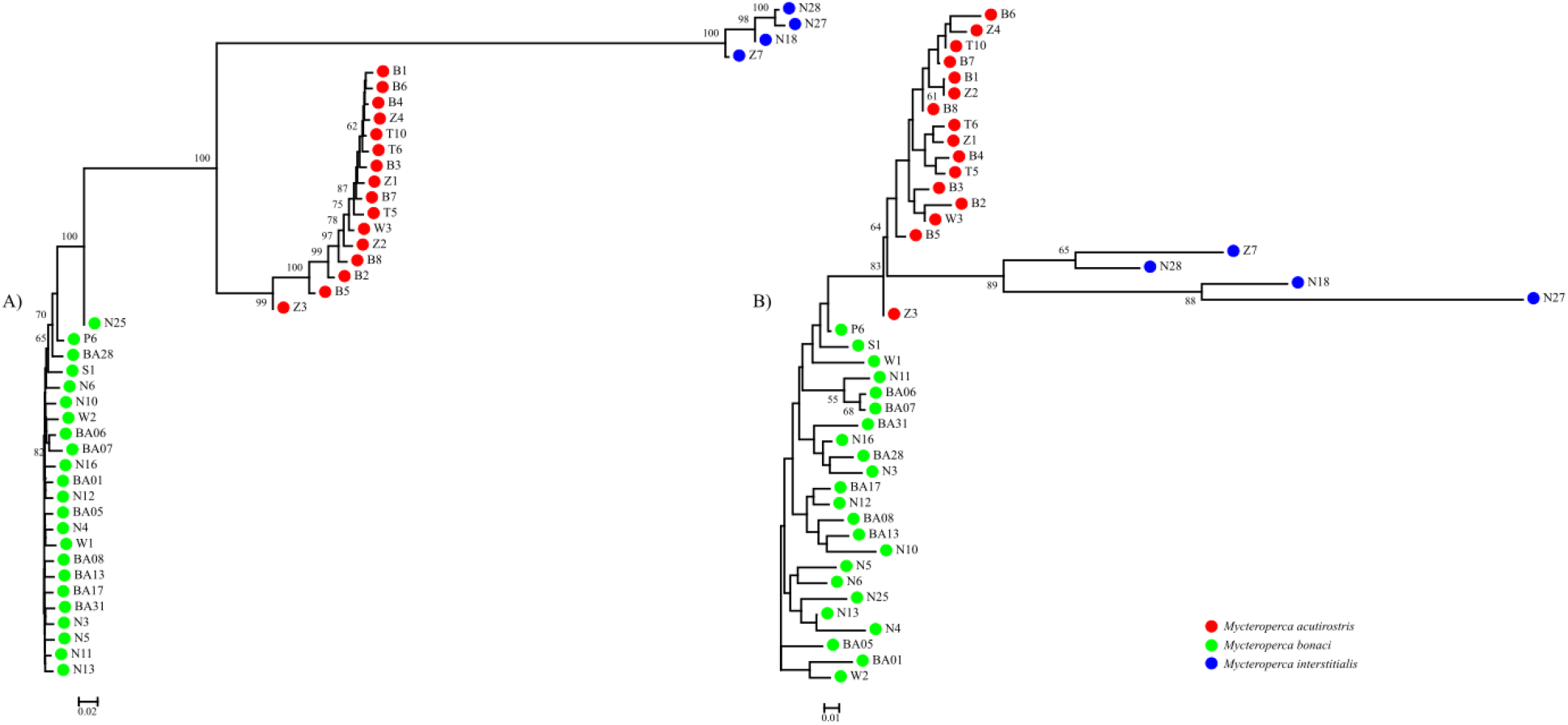
Neighbor-joining Trees Based on Nei’s Genetic Distances using the Following Sets of Loci: (A) a panel of 1,275 putatively neutral SNPs and (B) a panel of 38 putatively adaptive SNPs. Branch nodes are denoted as the percentage of bootstrap support that was generated with 1,000 replicates. Collection codes correspond to *M. acutirostis* (red dots), *M. bonaci* (green) and *M. interstitialis* (blue).

**Fig A2.**
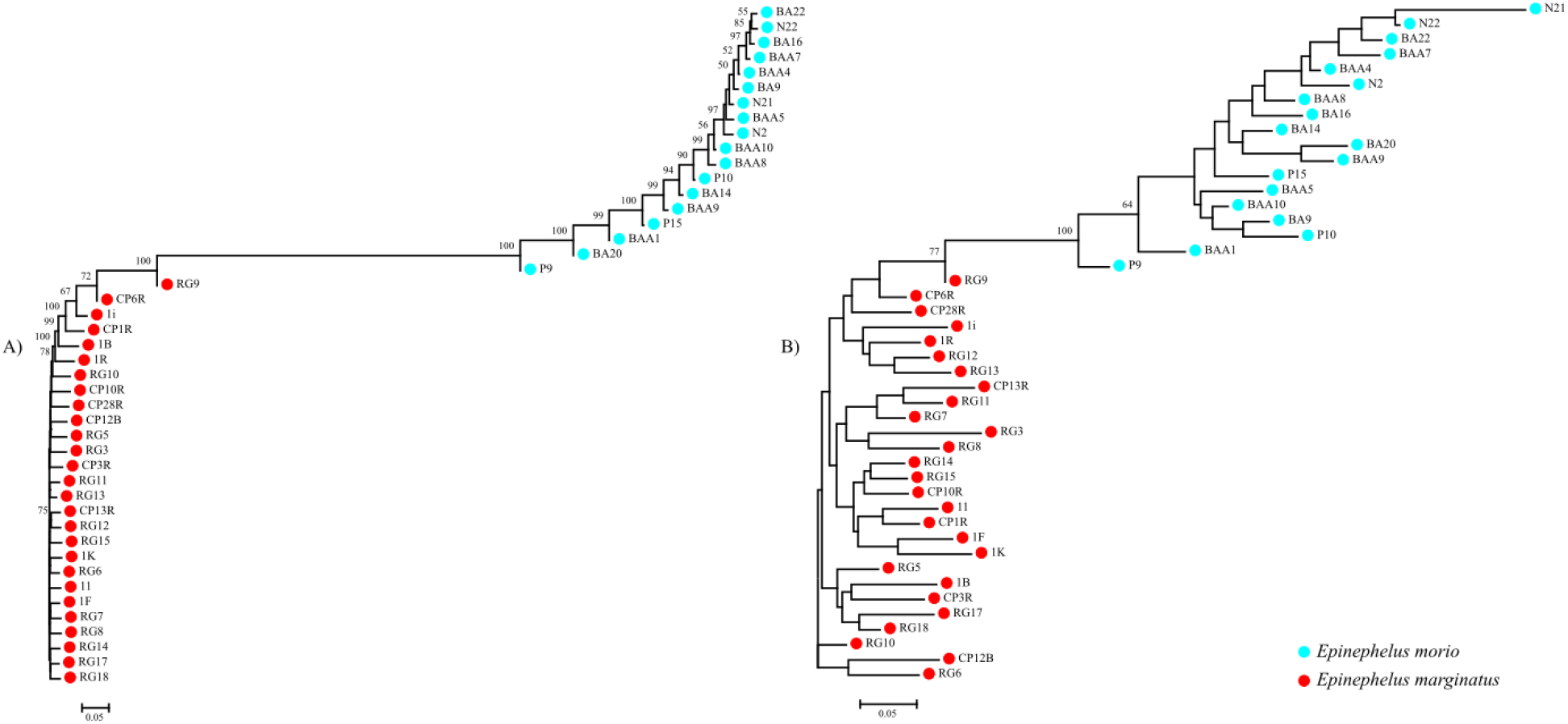
Neighbor-joining Trees Based on Nei’s Genetic Distances Using the Following Sets of Loci: (A) a panel of 3,490 putatively neutral SNPs and (B) a panel of 38 putatively adaptive SNPs. Branch nodes are denoted as the percentage of bootstrap support that was generated with 1,000 replicates. Collection codes correspond to *E. marginatus* (red dots) and *E. morio* (blue).

