## Supplementary figures and images for "Filling gaps: fishing, genetics, and conservation of groupers, especially the comb grouper (badejo) (*Mycteroperca acutirostris*), in SE Brazil (2013-2020)"

### Supplemental Figure 1

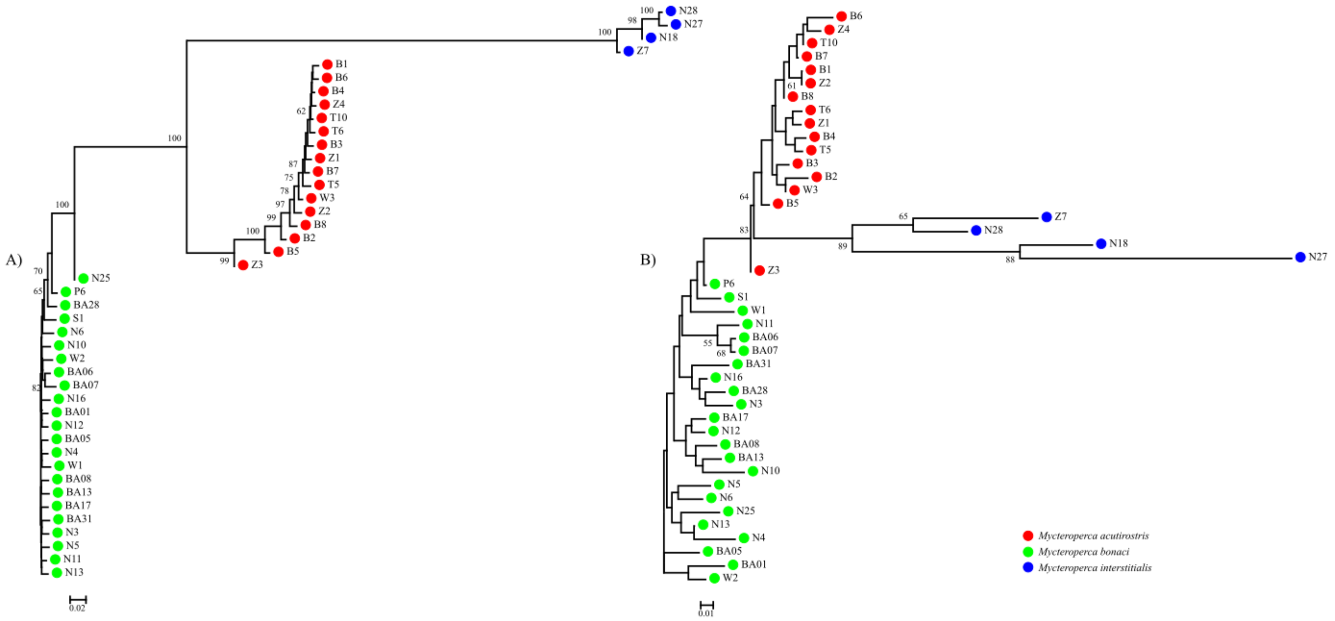

### Supplemental Figure 2

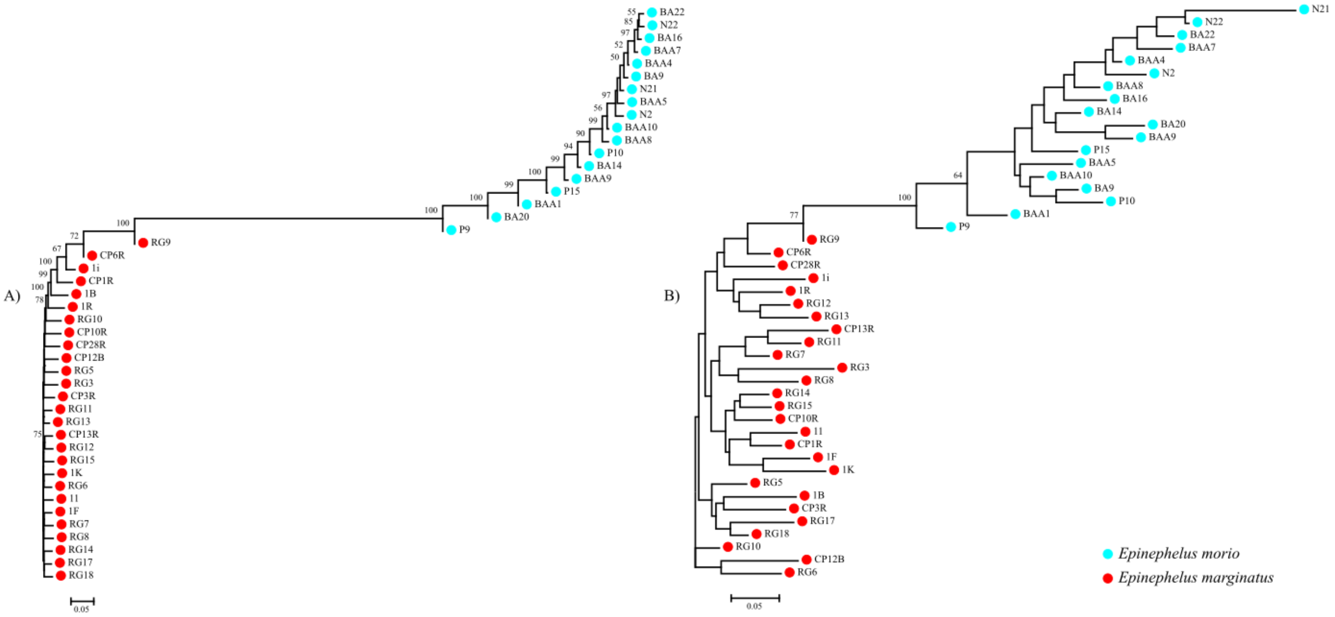
